# Small, Open-Source Text-Embedding Models as Substitutes to OpenAI Models for Gene Analysis

**DOI:** 10.1101/2025.02.15.638462

**Authors:** Dailin Gan, Jun Li

## Abstract

While foundation transformer-based models developed for gene expression data analysis can be costly to train and operate, a recent approach known as GenePT offers a low-cost and highly efficient alternative. GenePT utilizes OpenAI’s text-embedding function to encode background information, which is in textual form, about genes. However, the closed-source, online nature of OpenAI’s text-embedding service raises concerns regarding data privacy, among other issues. In this paper, we explore the possibility of replacing OpenAI’s models with open-source transformer-based text-embedding models. We identified ten models from Hugging Face that are small in size, easy to install, and light in computation. Across all four gene classification tasks we considered, some of these models have outperformed OpenAI’s, demonstrating their potential as viable, or even superior, alternatives. Additionally, we find that fine-tuning these models often does not lead to significant improvements in performance.

## Introduction

The integration of large language models (LLMs), particularly transformer models [1], into gene-expression data analysis has generated substantial interest due to their improved predictive capabilities across various applications (see, e.g., [2, 3, 4, 5, 6, 7, 8] for a review). Foundation models like scBERT [9], Geneformer [10], scGPT [11], and scFoundation [12] are trained on extensive collections of single-cell RNA-seq data that include thousands of datasets and millions of cells. This training process is designed to capture intricate information about genes and their interactions. Subsequently, these models are fine-tuned on much smaller, user-specific datasets tailored to specific applications. However, the development of foundation models from scratch necessitates significant resources. The extensive data collection and computational demands often make this process prohibitively expensive for most research groups. Additionally, deploying and fine-tuning these models for personalized applications presents challenges related to hardware requirements, software compatibility, and programming expertise [13].

The recently developed GenePT [14] introduces an innovative approach that eliminates the necessity of training foundation models from scratch. Utilizing OpenAI’s text-embedding function, this method transforms standard NCBI gene descriptions [15] into embeddings—dense numeric vectors that effectively encapsulate the textual content [16]. These embeddings comprehensively capture gene functions and their interactions as detailed in NCBI descriptions, offering a novel way to summarize and utilize existing knowledge about genes. These embeddings are directly employed as inputs for various machine learning methods, such as logistic regression and random forests, across diverse applications, such as classifying genes into drug-sensitive or drug-insensitive categories. The GenePT approach has been shown to achieve accuracy comparable to, or even surpassing, that of gene-expression foundation models in various gene-classification tasks [14].

The advantages of the GenePT approach are significant. It leverages OpenAI’s publicly accessible text-embedding function, eliminating the need for resource-intensive training of gene-expression foundation models. Additionally, this method is highly generalizable and can be extended to numerous other applications; OpenAI’s tool is capable of embedding descriptions of various biological entities, not limited to genes. Consequently, this approach is poised for broad adoption in the near future. Indeed, text-embedding plays a crucial role in natural language processing, where it is widely used in key applications such as text retrieval and translation, among others (see, e.g., [17] for a review).

However, relying on OpenAI’s text-embedding can also impose limitations on GenePT’s applicability. OpenAI’s embedding is an online service that requires users to send text to the OpenAI server and retrieve the embeddings from it. This reliance not only raises concerns about internet connectivity and security but also, and more critically, about the privacy of the uploaded text. While privacy might not be a significant concern for NCBI’s descriptions of genes, it becomes crucial if researchers wish to include their own insights, research results, or descriptions of other entities that are intended to remain confidential. The privacy issues associated with online LLMs have gained significant attention (see, e.g., [18, 19]); a recent study [20] reports that “all the methods leak query data,” and concludes that “to achieve truly privacy-preserving LLM adaptations that yield high performance and more privacy at lower costs, taking into account current methods and models, one should use open LLMs.” Additionally, using an online service raises other potential concerns, such as the reproducibility of results [21]. Furthermore, OpenAI charges a per-token fee for text embedding.

In this paper, we aim to address these limitations by replacing OpenAI’s service with open-source, transformer-based text-embedding models that can be installed and run locally. To ensure manageability for regular users with limited computational resources, we limit the size of these models to less than 100 million parameters. We refer to these as “small LLMs,” or S-LLMs for short. It is important to note that S-LLMs are text-embedding models designed to convert natural-language text into embeddings, and they are not the gene-expression foundation models discussed earlier. We will test these S-LLMs across the four gene-level applications discussed in the GenePT paper to determine if they perform comparably to GenePT’s approach, which relies on OpenAI’s embedding model. Specifically, we will employ logistic regression or random forest on the embeddings generated by S-LLMs or OpenAI’s model, and compare the accuracy of the predictions.

In the broader context of natural language processing, the performance of various embedding tools has been systematically assessed through the Massive Text Embedding Benchmark (MTEB) [22], which includes 58 datasets across 112 languages from eight embedding tasks, such as document classification, clustering, and retrieval. On average across these datasets, OpenAI’s embedding tool outperforms most open-source alternatives. However, some open-source models can perform just as well, or even better than, OpenAI’s tools in specific tasks. Therefore, it will be interesting to see whether the S-LLMs we have selected can match the performance of OpenAI’s embeddings in gene analysis.

If the S-LLMs cannot compete, there is a potential remedy: fine-tuning the S-LLMs. Specifically, instead of using the static text embeddings provided by the S-LLM as inputs for a separate classifier such as logistic regression or random forest, we could integrate an additional layer into the S-LLM, transforming it into a combined embedding and classification system. This integrated system would then be fine-tuned as a whole, allowing the embeddings to evolve during training. It is important to note that this fine-tuning process involves only minor adjustments to the weights in the pre-trained transformer model, resulting in a computational burden significantly lower than that of training an LLM from scratch. It is also worth noting that the capability to fine-tune is not available with OpenAI’s embedding function, which is closed-source online service.

## Results

### Settings of the comparisons

Our comparisons focus on the four gene-level tasks considered in the GenePT paper [23]. For the remainder of this paper, we will refer to these tasks as Task 1 through Task 4, respectively. Each of these tasks is a two-class classification problem, classifying genes into categories such as dosage-sensitive or dosage-insensitive transcription factors. Table 1 details the two classes and the sample size for each class. These datasets were curated, pre-processed, and made available by [10].

**Table 1:** Datasets and the number of samples in each class.

|  | Class 1 | Sample Size<br>in Class 1 | Class 2 | Sample Size<br>in Class 2 |
| --- | --- | --- | --- | --- |
| **Task 1** | Long-range<br>transcription factors | 46 | Short-range<br>transcription factors | 130 |
| **Task 2** | Dosage-sensitive<br>transcription factors | 122 | Dosage-insensitive<br>transcription factors | 368 |
| **Task 3** | Bivalent genes | 107 | Lys4-only-methylated<br>genes | 80 |
| **Task 4** | Bivalent genes | 107 | Non-methylated<br>genes | 42 |

For the gene descriptions, we utilized all human genes from the genome assembly version GRCh38.111, obtained from the `Ensembl` database [24]. Following the procedures outlined in the GenePT paper [23], we extracted descriptions from the summary section of the NCBI Gene database, after removing hyperlinks and date information. These cleaned descriptions were then used as input to generate text embeddings.

Hugging Face [25] offers a wide range of publicly available transformer-based text embedding models. These models vary greatly in size, measured by the number of parameters, and are categorized into five groups: fewer than 100 million, 100 million to 250 million, 250 million to 500 million, 500 million to 1 billion, and over 1 billion. Models on Hugging Face are systematically evaluated using the MTEB.

We selected ten models with fewer than 100 million parameters that ranked highest on the Hugging Face Leaderboard (https://huggingface.co/spaces/mteb/leaderboard_legacy, as of December 9th, 2024) for the “classification” embedding task in English. The models chosen include GIST-small-Embedding-v0 [26], NoInstruct-small-Embedding-v0 [27], stella-base-en-v2 [28], bge-small-en-v1.5 [29], e5-small-v2 [30], GIST-all-MiniLM-L6-v2 [26], gte-small [31], MedEmbed-small-v0.1 [32], e5-small [30], and gte-tiny [33]. A summary of these ten models is presented in Table 2, which includes the number of parameters, memory usage, average performance on MTEB classification datasets, and the embedding dimension each model produces. Notably, all of these S-LLMs contain no more than 55 million parameters. For context, OpenAI’s GPT-3 model contains 175 billion parameters—over a thousand times larger than our S-LLMs—and GPT-4 has 1.76 trillion parameters. These S-LLMs require a minimal amount of memory and can be easily installed and operated on a regular laptop computer. Except for stella-base-en-v2, which produces 768-dimensional embeddings, all other S-LLMs generate 384-dimensional embeddings.

**Table 2:** Information about the ten S-LLMs we use; more detailed information for each model can be found on the MTEB Hugging Face Leaderboard (https://huggingface.co/spaces/mteb/leaderboard_legacy).

| Models | Model parameters<br>(million) | Memory Usage<br>(GB) | MTEB Average<br>Performance | Embedding<br>dimensions |
| --- | --- | --- | --- | --- |
| GIST-small-Embedding-v0 | 33 | 0.12 | 76.1 | 384 |
| NoInstruct-small-Embedding-v0 | 33 | 0.12 | 76.0 | 384 |
| stella-base-en-v2 | 55 | 0.20 | 75.3 | 768 |
| e5-small-v2 | 33 | 0.12 | 72.9 | 384 |
| GIST-all-MiniLM-L6-v2 | 23 | 0.08 | 72.7 | 384 |
| gte-small | 33 | 0.12 | 72.3 | 384 |
| bge-small-en-v1.5 | 33 | 0.12 | 72.2 | 384 |
| MedEmbed-small-v0 | 33 | 0.12 | 71.8 | 384 |
| gte-tiny | 23 | 0.08 | 70.4 | 384 |
| e5-small | 33 | 0.12 | 68.1 | 384 |

For GenePT, we downloaded the gene text embeddings from its Zenodo repository. These embeddings were generated using OpenAI’s `text-embedding-ada-002` model [34], with each embedding having a dimension of 1536. This dimensionality is significantly higher than that of the embeddings produced by the S-LLMs.

### Logistic regression and random forests on S-LLM embeddings

For each of the four two-class classification tasks, we supplied gene embeddings generated by the ten S-LLMs, as well as those from OpenAI, to a classifier. We measured performance using the area under the ROC curve (AUC), which ranges from 0 to 1. Higher AUC values indicate better performance.

Following the GenePT paper, we chose logistic regression and random forest as our classifiers. These are implemented using the `LogisticRegression()` function from the `sklearn.linear_model` Python package and the `RandomForestClassifier()` function from the `sklearn.ensemble` Python package, respectively. We used the default hyperparameter settings for both functions without any further tuning.

For each task, we randomly split the data into training and testing sets, comprising 90% and 10% of the total data, respectively. The classifier—either logistic regression or random forest—is trained on the training data and then tested on the test data. To minimize the randomness in the evaluation, this entire procedure (random splitting, training, and testing) is repeated 10 times. Tables 3 and 4 present the mean AUC scores from these 10 repetitions, as well as the overall mean AUC scores across the four tasks. To aid in comparison, the tables also include the ranking (from highest to lowest AUC) of the embedding methods for each task and a final column that ranks the methods based on the overall mean AUC scores across all tasks, where a lower rank number indicates better performance.

**Table 3:** Performance of logistic regression with default hyperparameter settings on different embeddings.

| Embedding | AUC\|Rank<br>on Task 1 | AUC\|Rank<br>on Task 2 | AUC\|Rank<br>on Task 3 | AUC\|Rank<br>on Task 4 | Mean<br>AUC | Rank of<br>Mean AUC |
| --- | --- | --- | --- | --- | --- | --- |
| OpenAI | 0.712\|3 | 0.863\|11 | 0.931\|6 | 0.875\|11 | 0.845 | 7 |
| GIST-small-Embedding-v0 | 0.641\|7 | 0.887\|6 | 0.940\|1.5 | 0.917\|3 | 0.846 | 6 |
| NoInstruct-small-Embedding-v0 | 0.728\|2 | 0.908\|1 | 0.935\|3 | 0.907\|8 | 0.870 | 1 |
| stella-base-en-v2 | 0.702\|5 | 0.885\|8 | 0.934\|4 | 0.927\|1 | 0.862 | 4 |
| e5-small-v2 | 0.692\|6 | 0.879\|9 | 0.915\|10 | 0.922\|2 | 0.852 | 5 |
| GIST-all-MiniLM-L6-v2 | 0.608\|11 | 0.887\|5 | 0.940\|1.5 | 0.912\|5.5 | 0.837 | 8 |
| gte-small | 0.629\|8.5 | 0.866\|10 | 0.914\|11 | 0.915\|4 | 0.831 | 11 |
| bge-small-en-v1.5 | 0.629\|8.5 | 0.886\|7 | 0.924\|9 | 0.905\|9 | 0.836 | 9 |
| MedEmbed-small-v0 | 0.612\|10 | 0.892\|4 | 0.925\|8 | 0.910\|7 | 0.835 | 10 |
| gte-tiny | 0.706\|4 | 0.904\|2 | 0.930\|7 | 0.912\|5.5 | 0.863 | 3 |
| e5-small | 0.732\|1 | 0.898\|3 | 0.932\|5 | 0.892\|10 | 0.864 | 2 |

**Table 4:**
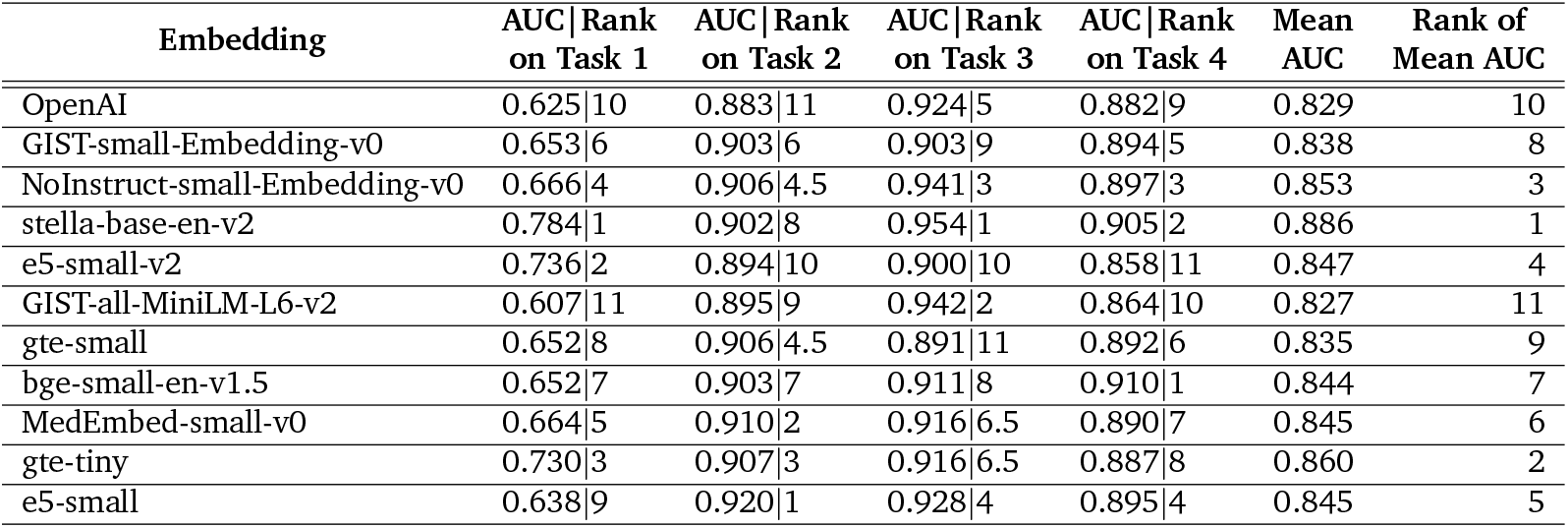
Performance of random forest with default hyperparameter settings on different embeddings.

| Embedding | AUC\|Rank<br>on Task 1 | AUC\|Rank<br>on Task 2 | AUC\|Rank<br>on Task 3 | AUC\|Rank<br>on Task 4 | Mean<br>AUC | Rank of<br>Mean AUC |
| --- | --- | --- | --- | --- | --- | --- |
| OpenAI | 0.625\|10 | 0.883\|11 | 0.924\|5 | 0.882\|9 | 0.829 | 10 |
| GIST-small-Embedding-v0 | 0.653\|6 | 0.903\|6 | 0.903\|9 | 0.894\|5 | 0.838 | 8 |
| NoInstruct-small-Embedding-v0 | 0.666\|4 | 0.906\|4.5 | 0.941\|3 | 0.897\|3 | 0.853 | 3 |
| stella-base-en-v2 | 0.784\|1 | 0.902\|8 | 0.954\|1 | 0.905\|2 | 0.886 | 1 |
| e5-small-v2 | 0.736\|2 | 0.894\|10 | 0.900\|10 | 0.858\|11 | 0.847 | 4 |
| GIST-all-MiniLM-L6-v2 | 0.607\|11 | 0.895\|9 | 0.942\|2 | 0.864\|10 | 0.827 | 11 |
| gte-small | 0.652\|8 | 0.906\|4.5 | 0.891\|11 | 0.892\|6 | 0.835 | 9 |
| bge-small-en-v1.5 | 0.652\|7 | 0.903\|7 | 0.911\|8 | 0.910\|1 | 0.844 | 7 |
| MedEmbed-small-v0 | 0.664\|5 | 0.910\|2 | 0.916\|6.5 | 0.890\|7 | 0.845 | 6 |
| gte-tiny | 0.730\|3 | 0.907\|3 | 0.916\|6.5 | 0.887\|8 | 0.860 | 2 |
| e5-small | 0.638\|9 | 0.920\|1 | 0.928\|4 | 0.895\|4 | 0.845 | 5 |

Surprisingly, OpenAI’s embedding does not emerge as the top performer on any of the four tasks, regardless of whether logistic regression or random forest is employed. With logistic regression, OpenAI’s embedding ranks 3rd, 11th, 6th, and 11th, with an overall ranking of 7th. Moreover, two of the S-LLMs we evaluated, NoInstruct-small-Embedding-v0 and e5-small, consistently outperformed OpenAI’s embedding across all four tasks. Similar results are observed with random forest: OpenAI’s embedding ranks 10th, 11th, 5th, and 9th, with an overall ranking of 10th. Additionally, three S-LLMs, NoInstruct-small-Embedding-v0, e5-small, and stella-base-en-v2, consistently performed better across all tasks.

### Logistic regression and random forests on S-LLM embeddings, with hyperparameter tuning

In the aforementioned analysis, we employed logistic regression and random forest classifiers without tuning their hyperparameters. For logistic regression, this involved using an *ℓ*_2_ penalty with a strength of 1, specified by setting `penalty = ‘l2’` and `C = 1.0` in the `LogisticRegression()` function. For random forest, we used a default configuration of 100 unpruned trees, set by `n_estimators = 100` and `max_depth = None` in the `RandomForestClassifier()` function.

The sub-par performance of OpenAI’s embedding observed in the previous section may be attributed to the lack of hyperparameter tuning. Therefore, in this section, we will tune these hyperparameters to see if the conclusions change. For logistic regression, we explored all combinations of `penalty = [‘l1’, ‘l2’]` and `C = [0.01, 0.1, 1, 10, 100]`. For random forest, we considered all combinations of `n_estimators = [25, 50, 100, 200, 400]` and `max_depth = [None, 10, 20, 30]`.

Once again, we randomly divided the data for each task into training and test datasets, consisting of 90% and 10% of all samples, respectively. We employed five-fold cross-validation on the training data to determine the optimal hyperparameters. These selected hyperparameters were then used to train a classifier on the entire training dataset. The classifier was subsequently tested on the test data, which had been set aside during the hyperparameter selection and training phases. This entire process was repeated ten times to minimize variability, and the mean AUC values were calculated and reported. The results for logistic regression and random forest are presented in Tables 5 and 6, respectively.

**Table 5:** Performance of logistic regression with hyperparameter tuning on different embeddings.

| Embedding | AUC\|Rank<br>on Task 1 | AUC\|Rank<br>on Task 2 | AUC\|Rank<br>on Task 3 | AUC\|Rank<br>on Task 4 | Mean AUC | Rank of<br>Mean AUC |
| --- | --- | --- | --- | --- | --- | --- |
| OpenAI | 0.674\|10 | 0.891\|11 | 0.934\|3 | 0.875\|11 | 0.843 | 10 |
| GIST-small-Embedding-v0 | 0.731\|3 | 0.897\|10 | 0.934\|3 | 0.903\|6 | 0.866 | 4 |
| NoInstruct-small-Embedding-v0 | 0.739\|2 | 0.905\|5 | 0.934\|3 | 0.910\|2 | 0.872 | 2 |
| stella-base-en-v2 | 0.723\|5 | 0.919\|2 | 0.915\|9 | 0.908\|3.5 | 0.866 | 3 |
| e5-small-v2 | 0.689\|9 | 0.912\|4 | 0.921\|6 | 0.918\|1 | 0.860 | 6 |
| GIST-all-MiniLM-L6-v2 | 0.622\|11 | 0.917\|3 | 0.943\|1 | 0.888\|8 | 0.842 | 11 |
| gte-small | 0.703\|7 | 0.903\|7 | 0.905\|11 | 0.908\|3.5 | 0.855 | 8 |
| bge-small-en-v1.5 | 0.700\|8 | 0.900\|9 | 0.911\|10 | 0.885\|9 | 0.849 | 9 |
| MedEmbed-small-v0 | 0.708\|6 | 0.901\|8 | 0.921\|6 | 0.893\|7 | 0.856 | 7 |
| gte-tiny | 0.725\|4 | 0.904\|6 | 0.919\|8 | 0.905\|5 | 0.863 | 5 |
| e5-small | 0.772\|1 | 0.928\|1 | 0.921\|6 | 0.883\|10 | 0.876 | 1 |

**Table 6:** Performance of random forest with hyperparameter tuning on different embeddings.

| Embedding | AUC\|Rank<br>on Task 1 | AUC\|Rank<br>on Task 2 | AUC\|Rank<br>on Task 3 | AUC\|Rank<br>on Task 4 | Mean<br>AUC | Rank of<br>Mean AUC |
| --- | --- | --- | --- | --- | --- | --- |
| OpenAI | 0.640\|7.5 | 0.883\|11 | 0.932\|4 | 0.880\|9 | 0.834 | 8 |
| GIST-small-Embedding-v0 | 0.682\|5 | 0.894\|10 | 0.934\|2.5 | 0.893\|7.5 | 0.851 | 5 |
| NoInstruct-small-Embedding-v0 | 0.657\|6 | 0.911\|2 | 0.914\|7 | 0.894\|6 | 0.844 | 7 |
| stella-base-en-v2 | 0.685\|4 | 0.908\|4 | 0.938\|1 | 0.900\|5 | 0.858 | 3 |
| e5-small-v2 | 0.740\|1 | 0.909\|3 | 0.908\|8.5 | 0.876\|10 | 0.858 | 2 |
| GIST-all-MiniLM-L6-v2 | 0.534\|11 | 0.900\|9 | 0.926\|6 | 0.871\|11 | 0.808 | 11 |
| gte-small | 0.602\|9 | 0.904\|8 | 0.906\|10 | 0.906\|2.5 | 0.829 | 10 |
| bge-small-en-v1.5 | 0.600\|10 | 0.908\|5 | 0.908\|8.5 | 0.919\|1 | 0.834 | 9 |
| MedEmbed-small-v0 | 0.640\|7.5 | 0.906\|7 | 0.931\|5 | 0.906\|2.5 | 0.846 | 6 |
| gte-tiny | 0.719\|2 | 0.908\|6 | 0.904\|11 | 0.893\|7.5 | 0.856 | 4 |
| e5-small | 0.715\|3 | 0.923\|1 | 0.934\|2.5 | 0.903\|4 | 0.869 | 1 |

Despite the adjustments, OpenAI’s embedding still does not outperform those from the S-LLMs, as detailed in the tables. It fails to achieve the highest AUC value in any of the four tasks, and on average across these tasks, it is ranked 10th and 8th out of 11 embeddings when using logistic regression and random forest, respectively. This means that for each task, at least one S-LLM embedding performs better than OpenAI’s. On average, nine out of ten S-LLMs outperform OpenAI when using logistic regression, and seven do so when using random forest.

Comparing Table 5 with Table 3, we observe that hyperparameter tuning enhances the AUC values for nine out of ten S-LLMs using logistic regression, with an average improvement of 0.011. In contrast, when using random forest, as shown by comparing Table 6 with Table 4, tuning improves AUC values for only four S-LLMs, with an average decrease of 0.003. This suggests that tuning the hyperparameters of the classifiers does not consistently enhance performance. This phenomenon is likely due to the small sample sizes of our tasks, coupled with high-dimensional input (the embeddings). For instance, in tasks 1 and 4, there are fewer than 50 samples in the minor class. This limited sample size, along with the high dimensionality of the input embeddings, contributes to significant randomness in the selection of optimal hyperparameters and high variance in the test AUC, especially for more complicated methods like random forest.

### S-LLM embeddings with fine-tuning

So far, we have demonstrated that the embeddings produced by S-LLMs often perform as well as, or even better than, those generated by OpenAI in our four tasks. These unexpected results render our initial plan to fine-tune S-LLMs to catch up with OpenAI’s performance unnecessary. However, we still wish to explore the fine-tuning of these S-LLMs in this section, with the aim of determining whether fine-tuning can further enhance their performance and thereby extend their advantage over OpenAI.

This fine-tuning approach differs significantly from the methods considered in previous sessions. Previously, all methods involved two independent steps. Initially, an LLM (either OpenAI’s or an S-LLM) would generate an embedding from text. Subsequently, a classifier (logistic regression or random forest) would use this embedding to predict the class label. These two steps were independent, meaning the embeddings generated in the first step did not change during the classifier’s training. Conversely, the fine-tuning approach employs a single integrated model. It involves adding a fully connected layer to the LLM, effectively transforming the LLM from an embedding generator into a classification model. This single deep-neural-network-based model is trained by taking NCBI gene descriptions as input and outputting the predicted class labels directly. During training, both the weights in the LLM and in the final layer are adjusted. Thus, the embeddings from the LLM are tuned specifically for the classification task, potentially enhancing classification accuracy.

For fine-tuning, we randomly divided the data for each task into training, evaluation, and test sets in proportions of 80%, 10%, and 10%, respectively. The fine-tuning process consisted of two stages. In the first stage, the model was trained on the training data and evaluated on the evaluation data to determine the optimal training hyperparameters. In the second stage, the model, configured with the best hyperparameters, was further trained on a combined set of the training and evaluation datasets, which accounted for 90% of all data, for several additional epochs to maximize the use of available information. Finally, this model was applied to the test data, which had been isolated from all fine-tuning and training activities, to compute the AUC score. This entire procedure was repeated ten times, and the mean AUC values were reported.

In the first stage of fine-tuning, we initialized the model with the following parameters: `learning_rate = 1e-5, num_train_epochs = 20, max_grad_norm = 0.7, warmup_ratio = 0.1`, and `weight_decay = 0.2`, with other parameters set to their default values. To prevent overfitting, we set up the early stopping `(EarlyStoppingCallback(early_stopping_patience = 5))` and the `ReduceLROnPlateau` scheduler `(factor = 0.8, patience = 2)` to gradually decrease the learning rate. In the second stage, the parameters were adjusted to `learning_rate = 1e-6, num_train_epochs = 10, max_grad_norm = 0.7`, and `weight_decay = 0.2`, with other settings remaining as default.

The computation was conducted on a single NVIDIA TITAN Xp GPU, an older model released in April 2017. The total time spent on the entire fine-tuning process, including data loading, model loading, and both stages of fine-tuning, is detailed in Table 7. For the two models with 23 million parameters, fine-tuning all four tasks takes about 20 minutes. Models with 33 million parameters take no more than 40 minutes, while the largest model, stella-base-en-v2, which has 55 million parameters, requires approximately one and a half hours. This demonstrates that the computational load is generally quite manageable, even on dated hardware.

**Table 7:** Computational time (in minutes) spent on fine-tuning S-LLMs under each task.

| Models | Time on<br>Task 1 | Time on<br>Task 2 | Time on<br>Task 3 | Time on<br>Task 4 | Mean<br>Time | Rank of<br>Mean Time |
| --- | --- | --- | --- | --- | --- | --- |
| GIST-small-Embedding-v0 | 33 | 36 | 32 | 30 | 32.8 | 6 |
| NoInstruct-small-Embedding-v0 | 26 | 33 | 26 | 24 | 27.3 | 3 |
| stella-base-en-v2 | 87 | 94 | 82 | 79 | 85.5 | 10 |
| e5-small-v2 | 32 | 35 | 31 | 32 | 32.5 | 5 |
| GIST-all-MiniLM-L6-v2 | 18 | 23 | 19 | 18 | 19.5 | 1 |
| gte-small | 36 | 43 | 37 | 32 | 37.0 | 8 |
| bge-small-en-v1.5 | 30 | 38 | 34 | 31 | 33.3 | 7 |
| MedEmbed-small-v0 | 37 | 44 | 37 | 39 | 39.3 | 9 |
| gte-tiny | 19 | 24 | 21 | 17 | 20.3 | 2 |
| e5-small | 26 | 31 | 29 | 25 | 27.8 | 4 |

Table 8 presents the results from fine-tuning the ten S-LLMs. Unexpectedly, compared to the results in previous tables, whether using logistic regression or random forest, with or without hyperparameter tuning, fine-tuning generally does not yield noticeable improvements in classification accuracy, if any. It is worth noting that we have tried more settings of training parameters than what we have described above. Specifically, we have tried `learning_rate = [1e-6, 5e-6, 1e-5, 5e-5, 1e-4], weight_decay = [0.0, 0.1, 0.2, 0.3, 0.4], warmup_ratio = [0.00, 0.05, 0.10, 0.15, 0.20], max_grad_norm = [0.6, 0.7, 0.8, 0.9, 1.0]`, and num_train_epochs = [5, 10, 15, 20, 25], and have not obtained results that are significantly better than those presented in Table 8.

**Table 8:**
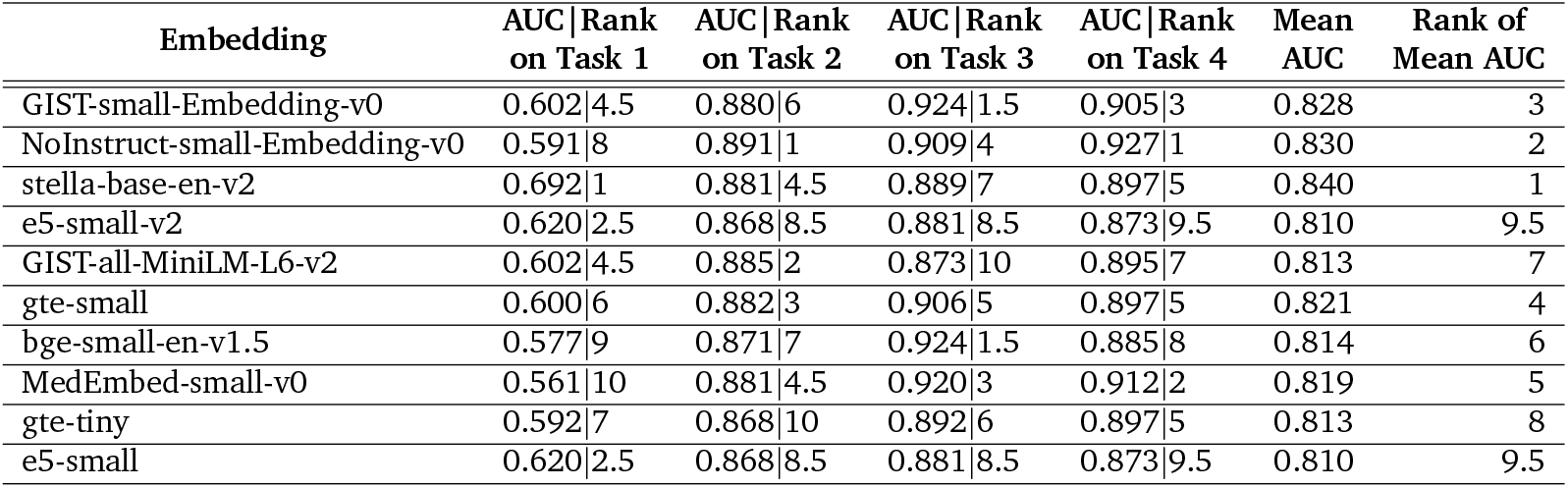
Performance of S-LLMs with fine tuning on different embeddings.

## Conclusions and Discussion

Recently, LLMs have been developed and utilized for describing and analyzing gene-expression data. While training a gene-expression foundation model can be labor-intensive and computationally prohibitive for most labs, GenePT’s approach of using gene embeddings generated by OpenAI’s text-embedding tools offers an affordable and efficient alternative. However, privacy concerns and other potential issues associated with the closed-source, online service nature of OpenAI’s embeddings may hinder the widespread adoption of GenePT’s approach. In this paper, we have explored the possibility of using open-source S-LLMs, which can be readily installed and run locally, as alternatives to OpenAI’s text-embedding. Our evaluation has focused on the four gene-level tasks considered in the GenePT paper.

While the expectation was that the embeddings from these S-LLMs could not compete with those from OpenAI, and that fine-tuning these S-LLMs might be necessary, the observations have been surprisingly different. We have found that even without fine-tuning, there are S-LLMs that outperform OpenAI on every task. Some S-LLMs even surpass OpenAI across all four tasks. This holds true regardless of whether logistic regression or random forest is used, and irrespective of whether the hyperparameters in the classifier are carefully tuned. This suggests that S-LLMs can be a legitimate alternative to OpenAI’s embedding tool for applications similar to those we have considered.

Second, we have found that although fine-tuning S-LLMs can be readily performed, it did not result in enhanced performance. This outcome is not entirely surprising. In fact, previous literature in more general settings of large language models, unrelated to genetics, has reported that fine-tuning pretrained LLMs is often brittle [35] and prone to degraded performance under small training sample sizes [36]. While our research does not preclude the possibility that fine-tuning might improve the performance of S-LLMs for gene analysis, and further investigation into this matter is certainly valuable, our findings suggest that simple, straightforward fine-tuning of S-LLMs may not consistently enhance their performance for gene analysis with small sample sizes.

Third, we have discovered that hyperparameter tuning slightly improves the performance of logistic regression but does not enhance the performance of random forest. Thus, using the default settings of these classifiers in Python functions can be a practical and reliable choice. Additionally, when comparing the AUC values generated by logistic regression with those produced by random forest, we observe that random forest often does not outperform logistic regression, despite its higher complexity and capability to capture non-linear relationships between features. These somewhat counterintuitive findings may be attributed to a couple of factors. First, the datasets we analyzed are characterized by high dimensions and small sample sizes, which makes reliable model selection very challenging. Second, the correlations or associations between the dimensions of the embeddings remain an open area of research, and it is still unclear whether random forest can effectively capture such relationships. Overall, using gene-description embeddings for gene analysis is a recent development. Further advancements in this field may hinge on a deeper understanding of these high-dimensional embeddings and how to utilize them most effectively. Additionally, exploring effective dimensionality reduction of these high-dimensional embeddings could also be a meaningful direction for future research.

Our current work has a limited scope, focusing primarily on gene-level analysis. GenePT also has potential applications at the cell level, such as cell-type clustering. As highlighted by [37], the question of how to efficiently combine gene embeddings with gene expression data to enhance the performance of gene-level tasks remains largely unanswered. Extending our analysis to cell-level tasks represents a challenging but potentially valuable future direction.

## Data availability

All datasets used in the study have been published and cited in the main body.

## Code availability

The source code is available at https://github.com/RavenGan/FinetuneEmbed.

## Acknowledgment

This work is supported by the National Institutes of Health (R01CA252878 to J.L.) and the DOD BCRP Breakthrough Award, Level 2 (W81XWH2110432 to J.L.).

